# Intersectional, anterograde transsynaptic targeting of the neurons receiving monosynaptic inputs from two upstream regions

**DOI:** 10.1101/2021.09.02.458803

**Authors:** Takuma Kitanishi, Mariko Tashiro, Naomi Kitanishi, Kenji Mizuseki

**Affiliations:** Department of Physiology, Osaka City University Graduate School of Medicine, Osaka 545-8585, Japan; PRESTO, Japan Science and Technology Agency (JST), Kawaguchi, Saitama 332-0012, Japan

**Keywords:** AAV1, anterograde transsynaptic tracing, INTRSECT, intersectional expression

## Abstract

A brain region typically receives inputs from multiple upstream areas. However, currently, no method is available to selectively access neurons that receive monosynaptic inputs from two upstream regions. Here, we devised a method to genetically label such neurons in mice by combining the anterograde transsynaptic spread of adeno-associated virus serotype 1 (AAV1) with intersectional gene expression. Injections of AAV1s expressing either Cre or Flpo recombinases and the Cre/Flpo double-dependent AAV into two upstream regions and the downstream region, respectively, were used to label the postsynaptic neurons receiving inputs from the two upstream regions. We demonstrated this labelling in two distinct circuits: the retina/primary visual cortex to the superior colliculus and the bilateral motor cortex to the dorsal striatum. Systemic delivery of the intersectional AAV allowed for unbiased detection of the labelled neurons throughout the brain. This strategy may help analyse the interregional integration of information in the brain.

## Introduction

The brain consists of many distinct regions with corresponding functions. A single brain region generally receives synaptic inputs from multiple upstream areas and distributes the information processed within the region to multiple downstream areas. Such integration and distribution are fundamental interregional interactions that support a variety of brain functions ^1,2^. The neurons that distribute information to two or more downstream areas through collateral projections can be readily labelled using multiple retrograde tracers ^3,4^. Moreover, these neurons can be genetically targeted and manipulated, and their activity can be monitored via several approaches, such as dual retrograde viral vector infection ^5^ and optogenetic identification of axonal projections during extracellular recordings ^6–9^. In contrast, although a range of viral vectors and transgenic animals has been successfully used to genetically target specific circuit structures ^10–12^, no method is currently available to selectively access the neurons that receive monosynaptic inputs from multiple upstream regions. Thus, in many brain regions, it remains unclear whether such neurons exist, and if so, what structures and functions are associated with them.

Here, we aimed to devise an intersectional, anterograde transsynaptic targeting that allows us to genetically access neurons that receive monosynaptic inputs from two upstream regions. This approach combines adeno-associated virus serotype 1 (AAV1) ^13,14^ with the previously developed intersectional expression system INTRSECT ^15,16^. High titers of AAV1 exhibit anterograde transsynaptic spread, which depends on neurotransmitter release machinery ^13,14^. By injecting the Cre recombinase-expressing AAV1 and the AAV with a Cre-inducible expression cassette into brain regions containing presynaptic and postsynaptic neurons, respectively, one can selectively label postsynaptic neurons receiving synaptic input from the presynaptic region. This technique has been widely applied in mice ^17–22^ and rats ^23–25^. INTRSECT is an intersectional gene expression system that depends on multiple recombinases ^15,16^. The C_on_/F_on_ expression cassette of the INTRSECT system turns on gene expression only when both Cre and Flpo recombinases are co-expressed in the target cells. We hypothesised that introducing the AAV1s expressing Cre and Flpo into two upstream brain regions and a C_on_/F_on_ expression cassette-containing AAV into the downstream region may aid in selectively labelling the neurons that receive monosynaptic inputs from both upstream regions (Fig. 1a). As a proof-of-principle, we aimed to demonstrate that postsynaptic neurons receiving both monosynaptic inputs can be selectively labelled in two distinct circuits: the retina/primary visual cortex (V1) to the superior colliculus (SC) and the bilateral secondary motor cortex (M2) to the dorsal striatum (DS) pathways. The labelling used here exhibited synaptic specificity with successful application in the two distinct circuits. We also demonstrated that the systemic delivery of the C_on_/F_on_ cassette is useful for the unbiased detection of neurons throughout the brain.

**Figure 1.**
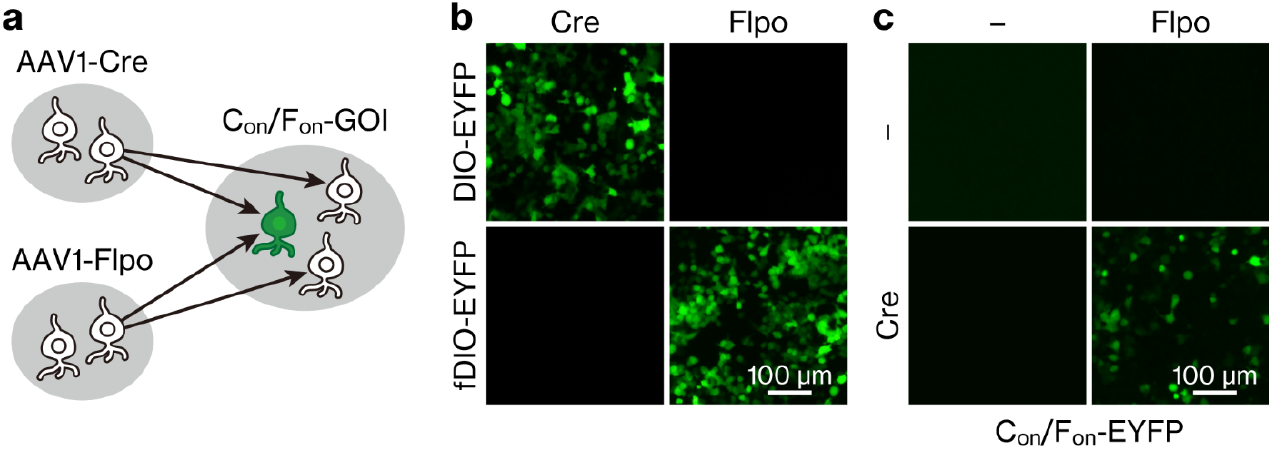
Concept of intersectional, anterograde transsynaptic targeting. **(a)** Schematic diagram of the intersectional, anterograde transsynaptic targeting of the neurons that receive monosynaptic inputs from two upstream brain regions. **(b)** EYFP fluorescence in HEK293T cells transfected with recombinase (pAAV-hSyn-Cre or pAAV-hSyn-Flpo-3×FLAG) and recombinase-dependent (pAAV-EF1α-DIO-EYFP or pAAV-EF1α-fDIO-EYFP) plasmid combinations. **(c)** EYFP fluorescence in HEK293T cells transfected with Cre, Flpo, and C_on_/F_on_ plasmid combinations. The pAAV-hSyn-C_on_/F_on_-EYFP plasmid was transfected into all wells. AAV1, adeno-associated virus serotype 1; GOI, gene of interest; EYFP, enhanced yellow fluorescent protein.

## Results

### Anterograde transneuronal labelling with AAV1-Cre and AAV1-Flpo

For the intersectional, anterograde transsynaptic targeting in the brain (Fig. 1a), we first confirmed the site-specific recombination using Cre and Flpo recombinases *in vitro*. HEK293T cells were transfected with recombinase-expressing plasmids (pAAV-hSyn-Cre or pAAV-hSyn-Flpo-3×FLAG) and recombinase-inducible, enhanced yellow fluorescent protein (EYFP)-expressing plasmids (pAAV-EF1α-DIO-EYFP or pAAV-EF1α-fDIO-EYFP). Cre/DIO and Flpo/fDIO combinations selectively induced EYFP expression (Fig. 1b), confirming that Cre and Flpo showed no cross-reactivity. For the C_on_/F_on_ plasmid (pAAV-hSyn-C_on_/F_on_-EYFP), EYFP was not expressed when only one of the Cre- or Flpo-expressing plasmids was transfected (Fig. 1c). EYFP was expressed only when both Cre- and Flpo-expressing plasmids were co-transfected (Fig. 1c), verifying the intersectional gene expression associated with the C_on_/F_on_ cassette ^15^.

We then tested the anterograde transneuronal spread of AAV1s expressing either Cre or Flpo under the hSyn promoter (hereafter, AAV1-Cre and AAV1-Flpo, respectively). The anterograde transneuronal spread with AAV1-Cre has been demonstrated in various glutamatergic and GABAergic pathways ^13,14,17–21,23–25^. However, it has only been demonstrated in a few pathways for AAV1-Flpo ^14,21,26^. In addition to the expected anterograde transneuronal spread of AAV1, it can also be transported in a retrograde direction from axons ^13,27,28^. This potential retrograde transport hampers the exclusive detection of cells labelled by anterograde transneuronal transport if a reciprocal connection exists between the targeted pre-and post-synaptic regions. Hence, we selected unidirectional pathways (V1 to ipsilateral SC, retina to contralateral SC, and M2 to contralateral DS) without back projection ^29–34^. We injected either AAV1-Cre and AAV1-EF1α-DIO-EYFP or AAV1-Flpo and AAV1-EF1α-fDIO-EYFP locally into the presynaptic and postsynaptic regions, respectively (Fig. 2a). In all three tested pathways, EYFP-positive somata were densely observed in the postsynaptic regions (i.e. SC or DS) for both AAV1-Cre and AAV1-Flpo (Figs. 2b to 2g). EYFP expression in the SC and DS was absent when either the AAV1-Cre or AAV1-Flpo injection to the presynaptic region was omitted (n = 4 mice), confirming the recombinase-dependent EYFP expression. These results suggested that both AAV1-Cre and AAV1-Flpo exhibited anterograde transneuronal spread capable of inducing site-specific recombination in postsynaptic neurons.

**Figure 2.**
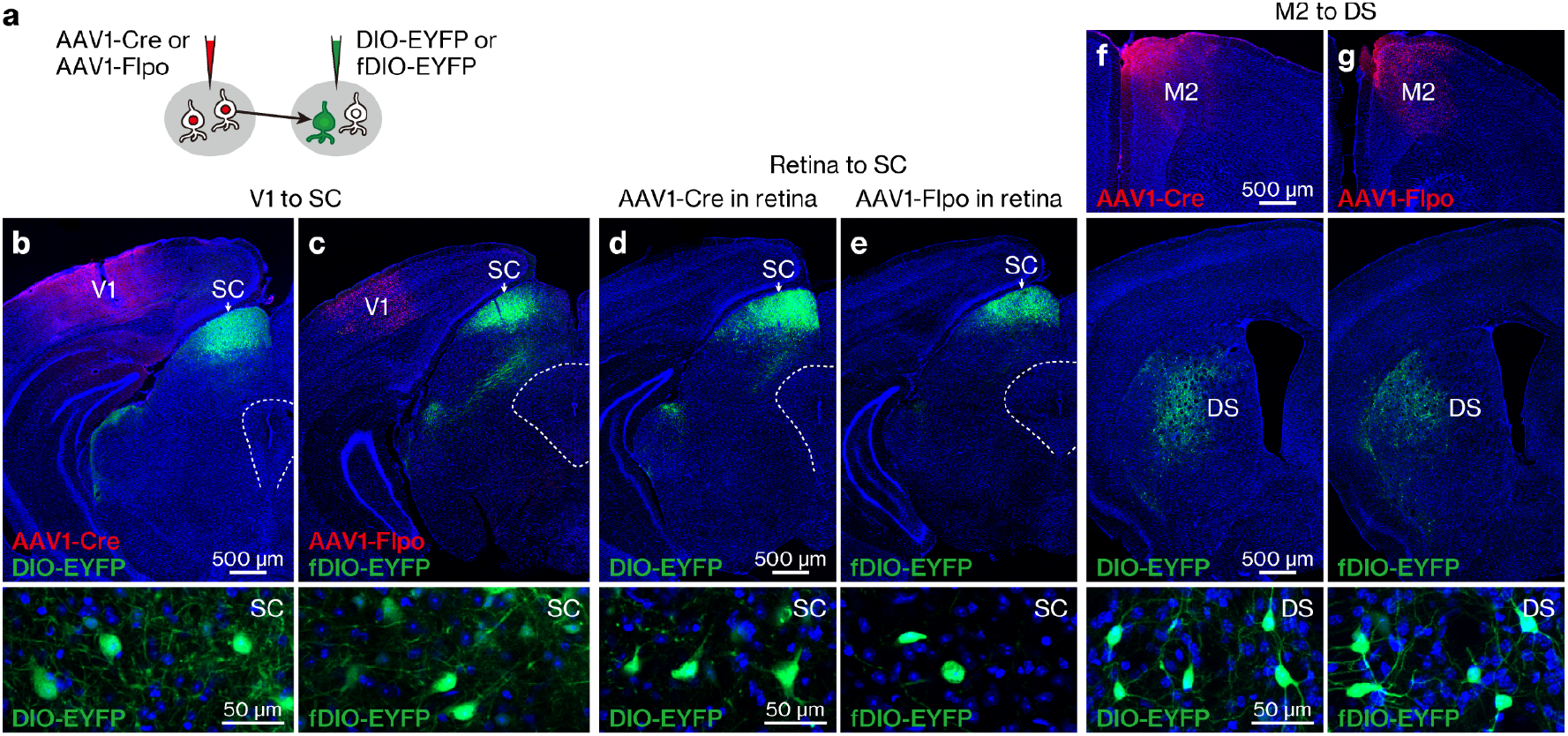
Anterograde transneuronal spread of AAV1-Cre and AAV1-Flpo. (**a**) Schematic diagram of the anterograde transneuronal labelling. (**b-g**) Anterograde transneuronal spread of AAV1-Cre (**b, d, f**) and AAV1-Flpo (**c, e, g**) from the V1 to ipsilateral SC (**b, c**), the retina to contralateral SC (**d, e**), and the M2 to contralateral DS (**f, g**) pathways. The Cre (red) and 3×FLAG-tagged Flpo (red) were detected with anti-Cre and anti-FLAG antibodies, respectively. All images represent coronal sections, and the bottom images show the labelled EYFP (green)-positive somata in the SC (**b-e**) and DS (**f, g**). Blue, DAPI. N = 2 (**b-e, g**) and 3 (**f**) results observed in mice under each experimental condition. AAV1, adeno-associated virus serotype 1; EYFP, enhanced yellow fluorescent protein; V1, primary visual cortex; SC, superior colliculus; M2, secondary motor cortex, DS, dorsal striatum.

### Intersectional, anterograde transsynaptic labelling in the retina/V1 to superficial SC (sSC) pathway

To genetically target neurons that receive monosynaptic inputs from two upstream regions, we performed intersectional, anterograde transneuronal labelling. The sSC receives synaptic inputs from both the retina and V1 ^35,36^, but whether individual sSC neurons receive monosynaptic inputs from both regions remains elusive ^32,37^. We injected AAV1-Cre, AAV1-Flpo, and AAV1-hSyn-C_on_/F_on_-EYFP into the right eye, left V1, and left SC, respectively (Fig. 3a). Together with Cre and Flpo expression in the injected regions (Figs. 3b and 3c), many EYFP-positive neurons were observed in the left sSC but not in the adjacent regions (Figs. 3c and 3d). EYFP expression was absent when either AAV1-Cre or AAV1-Flpo injection was omitted (n = 4 mice each), indicating that both AAV1-Cre and AAV1-Flpo, which were introduced to the presynaptic regions (i.e. retina and V1), were necessary for EYFP labelling in the downstream area (i.e. sSC).

**Figure 3.**
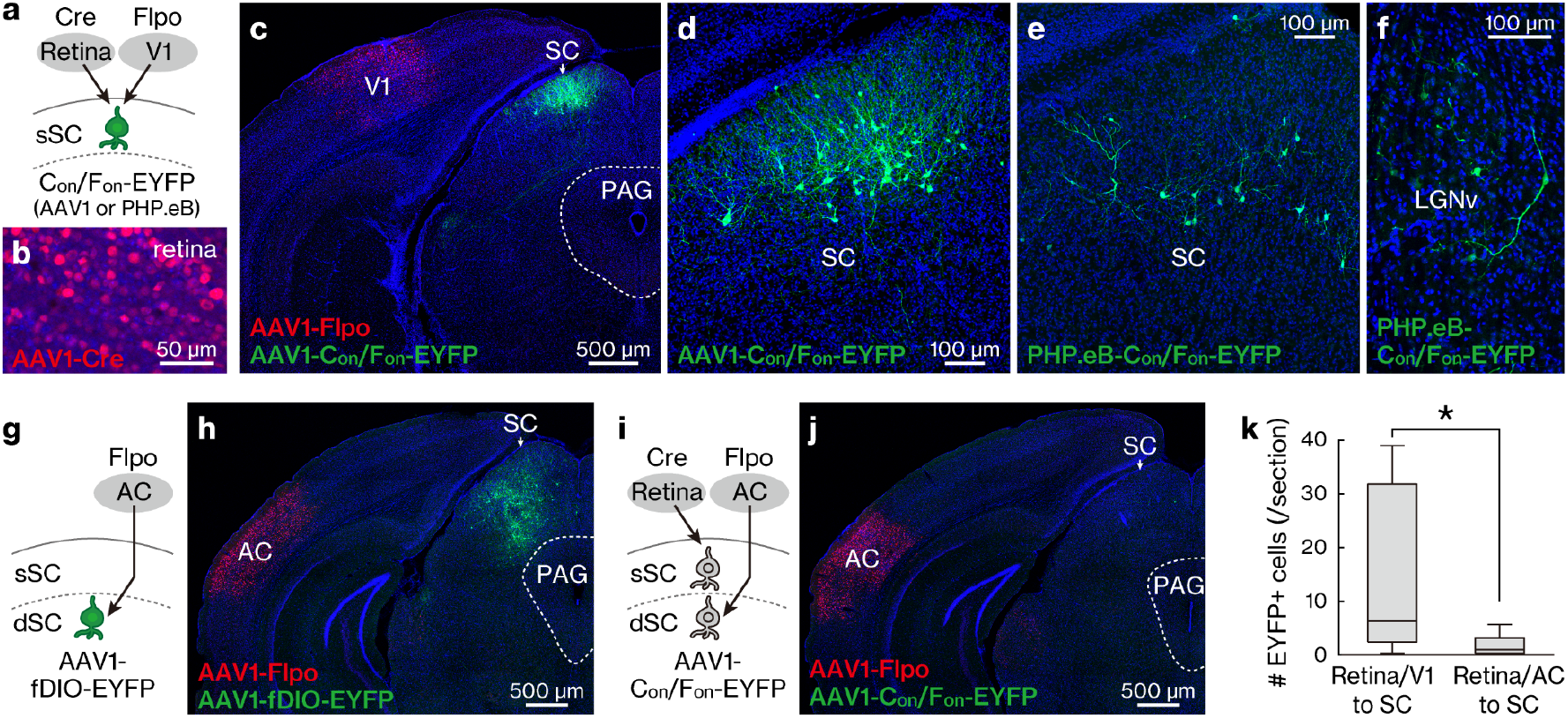
Intersectional, anterograde transsynaptic labelling of the retina/V1 to sSC pathway. (**a-f**) Intersectional, anterograde transsynaptic labelling of the retina/V1 to SC pathway. (**a**) Schematic diagram of the experiment. (**b**) Representative Cre (red) expression in the retina infected with AAV1-Cre. (**b-f, h, j**) Blue, DAPI. (**c-d**) Coronal sections showing the Flpo expression (**c**, red) in the V1 and the EYFP (green)-positive cells in the SC (**c-d**). The AAV1 containing C_on_/F_on_-EYFP expression cassette was injected locally in the SC. Number of mice = 7. (**d**) Magnified view of (**c**). (**e-f**) Coronal sections showing the EYFP (green)-positive cells in the left SC (**e**) and left LGNv (**f**). The C_on_/F_on_-EYFP expression cassette was delivered to the whole brain by the retroorbital injection of PHP.eB. N = 4 mice. (**g, h**) Anterograde transsynaptic labelling of the AC to deep SC (dSC) pathway. (**g**) Schematic diagram of the experiment. (**h**) Coronal section showing the EYFP (green) expression in the deep SC following the AAV1-Flpo (red) and AAV1-fDIO-EYFP injections into the AC and SC, respectively. N = 8 mice. (**i, j**) Lack of intersectional, anterograde transsynaptic labelling for the retina / AC to SC pathways. (**i**) Schematic diagram of the experiment. (**j**) Coronal section showing few EYFP (green)-positive cells in the SC following AAV1-Cre, AAV1-Flpo (red), and AAV1-C_on_/F_on_-EYFP injections into the retina, AC, and SC, respectively. N = 8 mice. (**k**) Numbers of EYFP-positive cells in the SC for the experiments of retina / V1 to SC (**c-d**) and retina / AC to SC (**j**) pathways. * *P* = 0.029, Mann-Whitney U test. Center line, median; box limits, upper and lower quartiles; whiskers, min and max. sSC, superficial superior colliculus; V1, primary visual cortex; LGNv, ventral lateral geniculate nucleus, EYFP, enhanced yellow fluorescent protein, AC; auditory cortex, AAV1, adeno-associated virus serotype 1.

While we delivered the C_on_/F_on_-EYFP expression cassette locally to the SC in the aforementioned experiment, delivering it to the whole brain would allow an unbiased detection of the inputs-integrating neurons throughout the brain. We tested this hypothesis using AAV-PHP.eB, which crosses the blood-brain barrier and transduces the whole brain following intravenous administration ^38^. We injected AAV1-Cre and AAV1-Flpo locally into the right eye and left V1, respectively, as well as AAV-PHP.eB-hSyn-C_on_/F_on_-EYFP intravenously via the retroorbital sinus. EYFP-positive cells were observed in the left sSC (Fig. 3e) and the left ventral lateral geniculate nucleus (LGNv) (Fig. 3f). The LGNv does not demonstrate any back projection to the retina or V1 ^29–32^, indicating that the observed EYFP-positive cells in the LGNv were likely the neurons integrating inputs from the retina and V1.

While neurons in the retina and V1 project to the sSC, those in the auditory cortex (AC) mainly project to the deep SC (dSC) ^35^. Taking advantage of these anatomically close but segregated pathways, we examined the synaptic specificity of the intersectional, anterograde transneuronal labelling. Injection of the AAV1-Flpo and AAV1-hSyn-fDIO-EYFP in the left AC and left SC resulted in preferential expression of EYFP in the left dSC (Fig. 3g and 3h), consistent with the known AC-to-dSC projection. Next, we injected AAV1-Cre, AAV1-Flpo, and AAV1-hSyn-C_on_/F_on_-EYFP in the right eye, left AC, and left SC, respectively (Fig. 3i). If AAV1 spreads selectively to mono-synaptically connected SC neurons, this injection would label almost no neurons in the SC, since the retina and AC neurons innervate mostly distinct SC layers with minor overlap ^30,39^. Following AAV injection, EYFP-positive somata were rarely observed in the SC (Fig. 3j), and their number was significantly smaller than that labelled in the retina/V1 to SC pathway experiment (Fig. 3k). This result suggested that anterograde transneuronal transport was selective for labelling expected postsynaptic SC neurons.

### Intersectional, anterograde transsynaptic labelling in the bilateral M2 to DS pathway

Next, we examined whether the intersectional, anterograde transsynaptic labelling can be applied to another circuit, the bilateral M2 to DS pathway. The DS receives synaptic inputs from both the left and right M2 with no back projection ^31,33,34^. The striatum is composed of medium spiny neurons, which are DARPP-32-positive principal cells comprising ~95% of the total striatal neuronal population, and interneurons including choline acetyltransferase (ChAT)-positive and parvalbumin (PV)-positive cells ^40^. All these types of neurons (i.e. medium spiny neurons, ChAT-positive cells, and PV-positive cells) receive monosynaptic inputs from M2 ^41–43^. However, it remains elusive whether, and if so, the type of individual DS neurons receive bilateral M2 monosynaptic inputs. To test this, we locally injected AAV1-Cre, AAV1-Flpo, and AAV1-hSyn-C_on_/F_on_-EYFP in the left M2, right M2, and bilateral DS, respectively (Fig. 4a). Cre and Flpo expression in the injected regions was confirmed by immunohistochemistry (Fig. 4b). In the DS, many EYFP-positive cells were observed bilaterally (Figs. 4c–4e). We found that the EYFP-labelled DS cells included DARPP-32-positive (mean ± s.d., 49.3 ± 9.6% of the EYFP-positive cells), ChAT-positive (16.1 ± 4.1%), and PV-positive (17.1 ± 2.3%) neurons (Figs. 4f to 4h). These results suggest that all types of DS neurons examined include cells that receive monosynaptic inputs from the bilateral M2.

**Figure 4.**
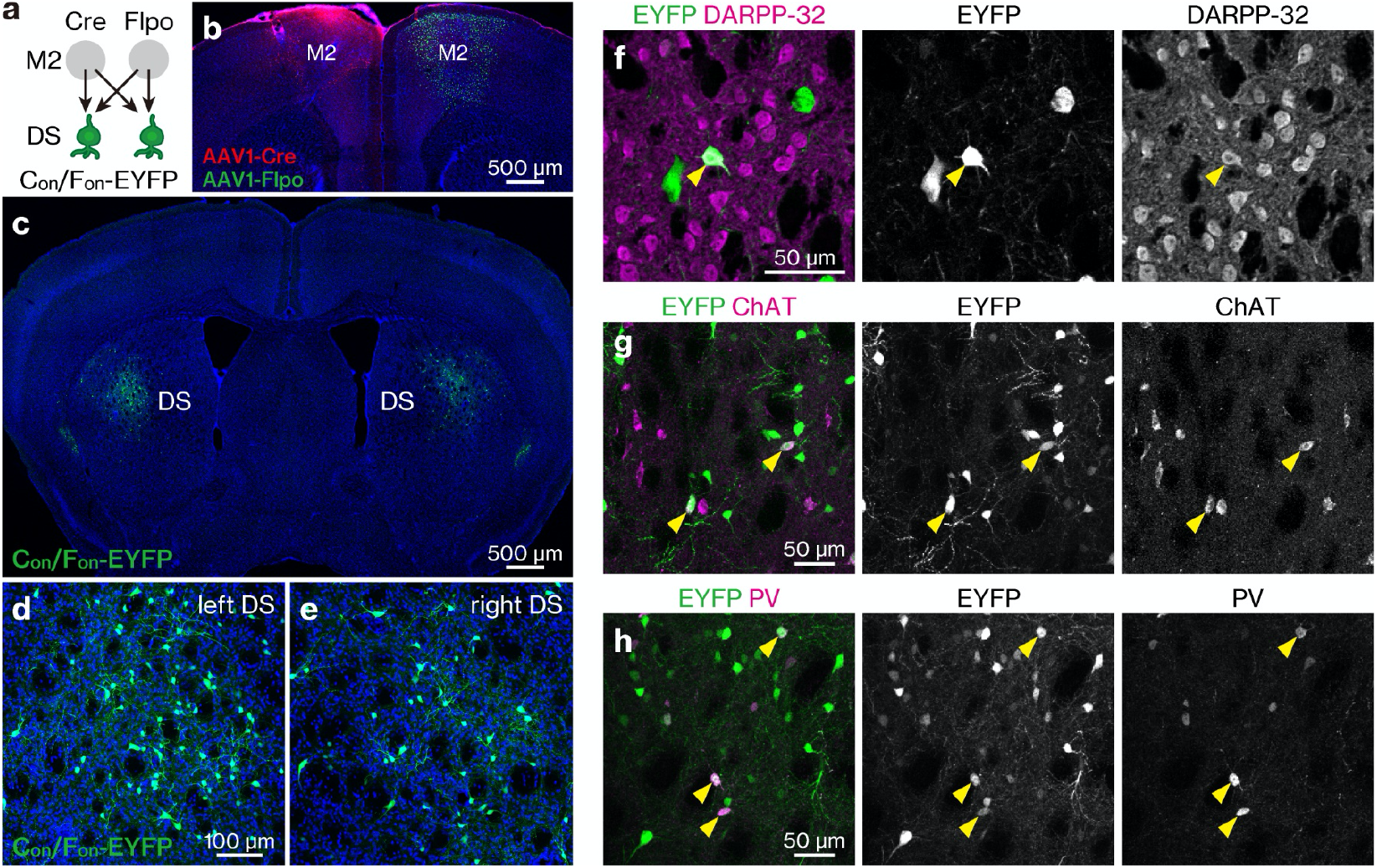
Intersectional, anterograde transsynaptic labelling of the bilateral M2 to DS pathway. (**a**) Schematic diagram of the experiment. (**b**) Cre (red) and Flpo (green) expressions in the left and right M2, respectively. (**b-e**) All images are coronal sections. Blue, DAPI. N = 4 mice. (**c-e**) EYFP (green) expression in the bilateral DS. (**d, e**) Magnified views of left (**d**) and right (**e**) DS. (**f-h**) Coronal DS sections showing the immunohistochemistry for dopamine- and cAMP-Regulated Phosphoprotein 32 kDa (DARPP-32) (**f**), choline acetyltransferase (ChAT) (**g**), or parvalbumin (PV) (**h**). Arrowheads, EYFP (green)-expressing cells double positive for the marker proteins (purple). M2, secondary motor cortex; DS, dorsal striatum; AAV1, adeno-associated virus serotype 1; EYFP, enhanced yellow fluorescent protein.

## Discussion

By combining the anterograde and transneuronal spread of AAV1 with intersectional gene expression, we established a method to genetically label neurons that receive monosynaptic inputs from two upstream regions. An anatomical control experiment suggested that this labelling exhibited synaptic specificity. We showed the success of this labelling method in two distinct circuits. We also demonstrated that the systemic delivery of the C_on_/F_on_ cassette is useful for the unbiased detection of neurons throughout the brain. Recently, the synaptic specificity of AAV1-based transneuronal labelling has been intensively characterised and established using anatomical, electrophysiological, and molecular approaches ^14^. In addition, AAV1 shows the transsynaptic spread in various pathways, including glutamatergic and GABAergic synapses ^14^. Taken together, our method provides a powerful means for determining the locations, numbers, and cell types of neurons receiving monosynaptic inputs from the two defined upstream regions.

Neurons that receive monosynaptic inputs from two upstream areas have been recently labelled using the combined introduction of AAV1-Cre, AAV1-Flp, Cre-dependent, and Flp-dependent expression cassettes in single mice ^14,26^. In these studies, the postsynaptic neurons of interest were identified as cells that co-express two fluorescent proteins with different colours. Instead of this approach, we used the C_on_/F_on_ intersectional expression cassette, which allowed us to specifically label the cells with a single gene of interest. This approach will be particularly useful for selectively manipulating and monitoring target cells with optogenetics, chemogenetics, imaging, and optogenetics-combined extracellular recordings ^10^. The intersectional expression system has been progressively expanded in several directions, including the use of mutually exclusive VCre recombinase ^16,44^, tetracycline-inducible expression system ^45^, and split-Cre complementation ^46^. Combining these systems with the present method may further enable us to target neurons that receive monosynaptic inputs from three or more upstream regions. Such intricately integrating ‘hub’ neurons might play a key role in generating novel information streams in neural circuits.

One of the limitations of AAV1-mediated transsynaptic tagging is that AAV1, similar to many other AAV serotypes, exhibits retrograde transport from the axons ^13,14,27,28,47,48^. Therefore, the application of this method is restricted to unidirectional pathways with no back projection ^13,14^. The mechanisms of directional transport remain unclear. However, the versatility of the AAV1-mediated transsynaptic tagging to dissect specific neurons in reciprocally connected circuits can be enhanced with significant improvements in the ratio of anterograde transsynaptic transport to retrograde transport via capsid engineering ^28,38^. Another limitation of this method involves the potential cytotoxicity of the Cre. Highly expressed Cre can damage cells by recognising DNA sequences that resemble *loxP* sites ^49–51^, whereas a relatively high AAV1 titer is necessary for its anterograde transsynaptic spread ^13^. This toxicity may be minimised by reducing Cre expression to a moderate level, for instance, by removing the woodchuck hepatitis virus post-transcriptional regulatory element (WPRE) from the AAV transfer plasmid while maintaining a high AAV1 titer^14^. Although these limitations need to be addressed, our method provides a straightforward approach to genetically dissect the neurons that directly receive multiple types of information.

## Methods

### Ethical declarations

All procedures related to animal care and use were approved by the Institutional Animal Care and Use Committee of Osaka City University (approved protocol #15030) and were performed in accordance with the *Guide for the Care and Use of Laboratory Animals* published by the National Institutes of Health.

### Plasmids

The following plasmids were obtained from Addgene: pAAV-hSyn-Cre-WPRE (no. 105553), pAAV-EF1α-mCherry-IRES-Flpo (no. 55634), pAAV-EF1α-DIO-EYFP (no. 20296), pAAV-EF1α-fDIO-EYFP (no. 55641), pAAV-hSyn-C_on_/F_on_-EYFP (no. 55650), and pUCmini-iCAP-PHP.eB (no. 103005). hSyn and EF1α denote the human synapsin 1 promoter and elongation factor 1 alpha promoter, respectively. The DIO, fDIO, and C_on_/F_on_ denote Cre-dependent, Flpo-dependent, and Cre/Flpo double-dependent gene expression cassettes, respectively ^15,52^. pXR1 was obtained from the National Gene Vector Biorepository ^53^. pHelper was obtained from the AAV Helper-Free System (no. 240071, Stratagene). During AAV1 preparation described below, we failed to obtain a high titer with pAAV-EF1α-mCherry-IRES-Flpo, which has a long insert (~5.0 kb) exceeding the AAV packaging limit of ~4.4 kb between the inverted terminal repeats (ITRs). Therefore, we constructed a shorter plasmid with a ~3.0 kb insert between ITRs, pAAV-hSyn-Flpo-3×FLAG (no. 173047, Addgene), which has the hSyn promoter followed by Flpo with 3×FLAG C-terminal tag (OGS629, Sigma-Aldrich). This plasmid was verified by sequencing and used throughout the study. Site-specific recombination of the DIO-, fDIO-, and C_on_/F_on_-EYFP cassettes by Cre and/or Flpo was confirmed by transfecting the corresponding plasmids (0.26 μg/well) into HEK293T cells (RCB2202, Riken BRC) seeded on 24-well tissue culture plates.

### AAV

AAV vectors were prepared as described previously with minor modifications ^25,48,54^. AAV serotype 1 (AAV1) was produced by co-transfecting three plasmids (pXR1, pHelper, and the pAAV plasmid containing the gene of interest; 20 μg/dish each) using PEI MAX (no. 24765-1, Polysciences) into HEK293T cells seeded on eight 15 cm tissue culture dishes. For AAV-PHP.eB^38^ production, the pXR1 plasmid was switched to pUCmini-iCAP-PHP.eB plasmid (20 μg/dish). The cells were harvested 72 h after transfection, and the AAV was purified using AAVpro Purification Kit Maxi (All Serotypes) (no. 6666, Takara Bio) according to the manufacturer’s instructions. The titer measurement via qPCR (StepOnePlus, Applied Biosystems) included the following: AAV1-Cre (1.8–2.3 × 10^13^ vg/ml), AAV1-Flpo (3.9 × 10^13^ vg/ml), AAV1-DIO-EYFP (1.4 × 10^13^ vg/ml), AAV1-fDIO-EYFP (1.3 × 10^13^ vg/ml), AAV1-C_on_/F_on_-EYFP (1.7 × 10^13^ vg/ml), and PHP.eB-C_on_/F_on_-EYFP (1.0 × 10^12^ vg/ml).

### Surgery

Three types of AAV injections were performed using male C57BL/6J mice (age on the day of the first surgery, 7.9–13.0 weeks; weight, 19.9–28.6 g; SLC, Japan) under anaesthesia (intraperitoneal injection of a mixture of 0.3, 4, and 5 mg/kg of medetomidine hydrochloride, midazolam, and butorphanol tartrate, respectively) ^55^. Intravitreal injection ^56^ was performed on the right eye to deliver AAV1-Cre to the retina. A small incision was made into the sclera using a 29-gauge needle (SS-05M2913, Terumo). A 33-gauge syringe (no. 80008, Hamilton) was inserted through the same incision to inject 1 μL of AAV1-Cre supplemented with 1:10 (v/v) 1% Fast Green FCF (no. 061-00031, Fujifilm Wako Chemicals) at a speed of 3–6 μL/min into the vitreous. The needle was kept inside the vitreous for 1–2 min before removal. Retroorbital injection ^57^ was performed for the systemic delivery of AAV-PHP.eB. A 29-gauge syringe (SS-05M2913, Terumo) was inserted into the right retroorbital sinus, and 10 μL of PHP.eB-C_on_/F_on_-EYFP (mixed with 40 μL of 1% Fast Green FCF and 50 μL of phosphate-buffered saline [PBS]) was slowly injected. Stereotaxic injections were performed to deliver AAV to specific brain areas. A craniotomy was performed above the injection site, and AAV was injected (5 μL/h) through a pulled glass pipette. Different AAVs were injected separately on different days with an interval of six or more days in the order of AAV1-Cre, AAV1-Flpo, and one of the DIO-, fDIO-, or C_on_/F_on_-EYFP. This multi-step stereotaxic injection was used to minimise any risk of contamination among the AAVs during surgery ^13,25^. The coordinates ^58^ and volumes are listed as follows: left V1 (anteroposterior from bregma [AP], –3.9 mm; mediolateral from the midline [ML], 2.6 mm; dorsoventral from the cortical surface [DV], 0.6 mm; 0.2–0.4 μl), left AC (AP, –3.1 mm; ML, 4.0 mm; DV, 0.75 mm; 0.2 μl), left SC (AP, –3.9 mm; ML, 0.8 mm; DV, 1.5 mm; 0.4 μl), M2 (AP, 1.5 mm; ML, 0.8 mm; DV, 0.8 mm; 0.2–0.4 μl/region), and DS (AP, 0.5 mm; ML, 2.3 mm; DV, 3.0 mm; 0.4 μl/region).

### Histology

Mice were transcardially perfused with 0.9% saline and then 4% paraformaldehyde in 0.1 M phosphate buffer under anaesthesia after 18 ± 4 days following the last surgery. The brains and the eyes (for mice with intravitreal injection) were enucleated and stored in the same fixative overnight at 4 °C. The brains were transferred and kept in 30% sucrose in PBS for more than 48 h at 4 °C and then sectioned using a freezing microtome (SM2010R, Leica; EF-13, Nihon Microtome Laboratory) at a thickness of 40 μm parallel to the coronal plane. Whole-mount retinae were prepared by removing the cornea, sclera, and lens from the eyes and by making radial incisions into the retina ^59^. The sections and retinae were incubated sequentially with 5% bovine serum albumin (BSA)/0.3% Triton X-100 in PBS for either 30 min (for sections) or 60 min (for retinae) at room temperature, primary antibodies in 5% BSA/PBS overnight at 4 °C, and corresponding secondary antibodies with DAPI (0.5 μg/mL, D1306, Thermo Fisher) in 5% BSA/PBS either for 2 h at room temperature or overnight at 4 °C. The samples were washed with PBS between the incubations. The following primary antibodies were used: mouse anti-Cre recombinase (1:2,000, MAB3120, Millipore), rabbit anti-FLAG (DYKDDDDK) (1:1,000, RFLG-45A-Z, ICL), chicken anti-GFP (1:2,000, ab13970, Abcam; only for sections with PHP.eB-C_on_/F_on_-EYFP), rabbit anti-DARPP-32 (1:800, 2306S, Cell Signaling Technology), goat anti-ChAT (1:100, AB144P, Millipore), and rabbit anti-PV (1:4,000, PV27, Swant). The following secondary antibodies (all from Thermo Fisher) were used at a dilution of 1:800: goat anti-chicken IgY conjugated with Alexa Fluor 488 (A-11039), goat anti-mouse IgG with Alexa Fluor 594 (A-11032), goat anti-rabbit IgG with Alexa Fluor 594 (A-11037), donkey anti-goat IgG with Alexa Fluor 594 (A-11058), and goat anti-rabbit IgG with Alexa Fluor 647 (A-21245). The sections and retinae were mounted on coverslips with an antifade mountant (P36961, Thermo Fisher). Fluorescent images were obtained using confocal microscopes (LSM800 and LSM700, Zeiss) equipped with 10× (numerical aperture = 0.45) and 20× (numerical aperture = 0.8) objectives. Cells positive for EYFP, ChAT, or PV were detected from the fluorescent images using automated software (Image-based Tool for Counting Nuclei, Center for Bio-image Informatics at University of California, Santa Barbara), followed by manual correction of erroneous detections ^25^. DARPP-32-positive cells were detected manually using Fiji software ^60^.

### Statistics and reproducibility

Statistical analysis was performed using IBM SPSS Statistics (ver. 21, IBM). *P* < 0.05 obtained using the Mann-Whitney U test was considered as a significant difference. All data are reported as the mean ± standard deviation unless otherwise specified.

## Acknowledgements

We thank the Research Support Platform, Osaka City University Graduate School of Medicine for assistance with confocal microscopy. This work was supported by JST PRESTO (JPMJPR1882 to T.K.), JSPS KAKENHI (20H03356 and 17K19462 to K.M. and 20K06878 and 19H04937 to T.K.), Takeda Science Foundation (to K.M.), The Uehara Memorial Foundation (to K.M. and T.K.), The Naito Foundation (to K.M. and T.K.), Kato Memorial Bioscience Foundation (to T.K.), and Osaka City University Strategic Research Grant for Young Researchers (to T.K.).

## Author contributions

T.K. and K.M. conceived the project. T.K., M.T., and N.K. performed the experiments. T.K. and K.M. wrote the manuscript with input from all the authors.

## Data availability

The pAAV-hSyn-Flpo-3×FLAG plasmid is available from Addgene (no. 173047). The data supporting this study are available from the corresponding authors upon reasonable request.

